# A novel high-throughput assay identifies small molecules with activity against persister cells

**DOI:** 10.1101/2023.04.13.536681

**Authors:** Maiken Engelbrecht Petersen, Liva Kjær Hansen, Nicholas M. Kelly, Thomas Keith Wood, Nis Pedersen Jørgensen, Lars Jørgen Østergaard, Rikke Louise Meyer

## Abstract

Persister cells are a subpopulation of transiently antibiotic tolerant bacteria, which are believed to be the main cause of relapsing bacterial infections. Due to the importance of persister cells in human infections, there is a need for new antibiotics that kill bacteria independently of their activity. However, high-throughput assays to screen for drugs with such activity are missing. This is partly due to the transient nature of the phenotype, which makes it is difficult to prepare a concentrated population of persister cells that remain inactive during incubation with antibiotics in standard growth media.

The purpose of this study was to develop a simple and high-throughput assay to identify compounds with antimicrobial activity against persister cells during a 24 h incubation period. Subsequently, this assay was used to screen a selection of small molecules with hypothesized antimicrobial activity.

The fraction of *S. aureus* that tolerate bactericidal concentrations of ciprofloxacin were defined as persister cells. We first quantified how the cell concentration, growth phase, antibiotic concentration, duration of antibiotic exposure, and presence/absence of nutrients during antibiotic exposure affected the fraction of persister cells in a population. After optimizing these parameters, we compared our approach to generate persister cells, to a process that generated persister cells by a short exposure to rifampicin. Finally, we used the optimised protocol to identify molecular structures that have anti-persister activity by performing screening on initially compound fragments and then selecting compounds that incorporated the fragments that displayed activity.

We show that exponential- and stationary-phase cultures transferred to nutrient-rich media only contain a small fraction (0.001 to 0.07 %) of persister cells that tolerated 10, 50 and 100 × MIC ciprofloxacin. Exponential-phase cultures displayed a bi-phasic time-kill curve, which plateaued after 5 h exposure, while stationary phase cultures displayed a low, but constant death rate at 50 and 100 × MIC ciprofloxacin. Inducing the persister phenotype with a short rifampicin treatment resulted in 100% persister cells when evaluated after ≤5 h exposure to ciprofloxacin. However, after longer incubation times, cells resumed activity and lost their tolerance to ciprofloxacin. Tolerance was only maintained in the majority of the population for the full 24 h incubation period if cells were transferred to a carbon-free minimal medium before exposure to ciprofloxacin. We conclude that keeping cells starved in a carbon-free medium enables generation of high concentrations of *S. aureus* cells that tolerate 50 × MIC ciprofloxacin, and we find this protocol easily applicable for rapid screening of anti-persister drugs that act on dormant or non-dividing cells.

## Introduction

Gladys Hobby first discovered bacterial persister cells in 1942 by observing a small percentage of surviving bacteria after treating a culture of cocci with penicillin^1^. She further observed that these survivors were non-growing. Two years later, Joseph Bigger named this tolerant sub-population and he discovered that the survivors where indistinguishable from the original culture^2^. He hypothesised that these persisters were no more resistant to penicillin than their fellows, but instead they were tolerant to the action of penicillin. Today, persistence is defined as the ability of a subset of a bacterial population to survive exposure to a bactericidal drug concentration - although they cannot replicate in the presence of the drug^3^. The presence of persister cells in a population is characterised by a bi-phasic time-kill curve, where susceptible cells are killed at an initial fast rate, leaving behind a population of persister cells that tolerate the antibiotic and are killed at a much slower rate^3^.

Persister cells are non-growing and have activated a suite of protective mechanisms, which allow them to temporarily cope with starvation or different forms of stress. These include cell envelope stress^4^, starvation^5^, oxidative stress^6,7^, heat stress, DNA damage, and accumulation of misfolded proteins^8^. By arresting growth, slowing metabolism, and activating protective mechanisms, persister cells tolerate extremely high concentrations of antibiotics, as antibiotics typically target processes involved in growth and cell division, or require a certain level of metabolic activity to exert their antimicrobial effect^9–11^. The persister phenotype is often associated with bacterial biofilms, where persister cells exist as a sub-population. Inside the biofilm, bacteria are protected from the immune system, and the non-growing persister cells are thus protected from elimination by professional phagocytes. The persister phenotype is reversible, and bacteria resume growth when conditions are favourable. Therefore, the persister phenotype is often associated with recalcitrant and relapsing bacterial infections^12,13^, and there is an urgent need for antibiotics that not only target actively growing bacteria, but also persister cells.

Efforts to discover novel antibiotics often rely on high-throughput assays to screen large compound libraries for antimicrobial activity. Standard assays detect antimicrobial activity based on inhibition of bacterial growth in a rich laboratory medium^14–16^. However, this approach is biased towards identifying compounds that target actively growing bacteria and will therefore most likely identify compounds with the same shortcomings as currently available antibiotics. Antibiotics that target persister cells must kill bacteria independent of their metabolic state. High-throughput screening assays designed for discovery of anti-persister drugs must therefore measure the loss of viable cells from a highly concentrated population of persister cells.

The first challenge for designing an anti-persister drug screening assay is to reproducibly generate a high concentration of persister cells. Ideally, assays that screen for anti-persister drugs should detect a 3-log reduction in colony forming units (CFU) to establish biocidal activity^17^. This seemingly simple requirement is difficult to meet, because persister cells comprise a small and rather variable fraction of bacterial populations. Experimentally, persister cells have been quantified by measuring surviving cells by CFU enumeration after challenging a population with high concentrations of antibiotics (≥10 × MIC) in a laboratory growth medium for some hours. The incubation time must be long enough to kill susceptible cells, such that viable cells are enumerated after the bi-phasic time-kill curve has reached the second phase with the slow death-rate representing persister cells^18^. Many studies have estimated the fraction of persister cells in a stationary phase broth culture after 3-5 h incubation with the antibiotic, and the number varies greatly from 0.000001% to just under 1%^18–21^. After such short incubation time, the susceptible population may not be completely eliminated (i.e. the population is still on the steep part of the time-kill curve), which results in highly variable and time-sensitive results^22^. A reproducible assay for identifying anti-persister drugs should therefore involve much longer incubation times, e.g. 24 h which standard in other antimicrobial assays.

The media composition, nutrient concentration, and exposure to different stress factors affects how many cells in a population will transition to the persister state. Furthermore, these and other unknown factors affect how quickly the bacteria resuscitate and become susceptible to antibiotics after transfer to fresh growth media. Since the persister fraction is quantified as the population of cells that survive exposure to antibiotics after a specified amount of time, the dynamics of resuscitation will also impact the fraction of viable cells present at the time of sampling. While the death rate of the susceptible population in the first phase of the time-kill curve reflects how fast the antibiotic acts on the cells, it has been shown that the death rate in the second phase does not reflect the action of the antibiotic on non-dividing cells, but rather reflects the rate at which a population of inactive cells resume activity and become susceptible^23^. It is thus difficult to discern if a drug is truly active against bacteria in the persister state when testing the efficacy of the drug in a nutrient-rich medium where cells can resume activity.

Much research on persister cells has been performed on *Escherichia coli*^20,23–25^, and Kwan et al. reported that *E. coli* persister formation could be triggered by treating exponential phase cultures briefly with rifampicin, tetracycline or carbonyl cyanide *m*-chlorophenylhydrazone (CCCP) to halt transcription, translation or ATP-synthesis, respectively. This approach resulted in persister fractions of 10-100% in *E. coli*^20^ as assessed by their tolerance to ciprofloxacin and ampicillin for 3 h. In another study, the pre-treatment using rifampicin initially killed approx. 66% of the population before turning almost all the remaining cells into highly tolerant persister cells^24^, as shown by their tolerance to ampicillin for 2 h. However, protocols for generating persister cells are less well characterised in staphylococci, which are the culprits of many chronic biofilm-associated infections^26^. For *Staphylococcus aureus*, one study showed that CCCP exposure induced the persister phenotype in 60% of the population, as shown by their tolerance to levofloxacin for 3 h^27^, indicating that ATP levels also play a role for Gram positive bacteria. The mechanism for persister formation presumably varies between different bacterial species, particularly between Gram positive and Gram negative bacteria. Knowledge is therefore not necessarily transferable between bacterial strains and growth conditions, and protocols for preparation of antibiotic-tolerant persister cells must therefore be validated for each experimental setting.

The aim of this study was to establish a high-throughput screening protocol to identify small molecules with antimicrobial activity against *S. aureus* persister cells. We investigate the use of rifampicin as a pre-treatment step to induce persister formation, and we characterize how the experimental conditions for antibiotic exposure affect the fraction of cells that tolerate antibiotics and can be perceived as persister cells. These experimental conditions include the bacterial cell concentration, the antibiotic concentration, the exposure time, and the presence of nutrients during antibiotic exposure. Finally, we use the optimised protocol to identify molecular structures that have anti-persister activity. Approximately 250 compounds from a fragment library were screened resulting in a cluster of fragments having anti-persister activity with the same common structural motif. Development of this fragment to improve activity and physiochemical properties utilising commercially available compounds incorporating was then undertaken.

Our objectives in designing an assay to screen for anti-persister drugs was therefore 1) to generate a high concentration of persister cells to enable detection of a 3-log reduction, and 2) to expose the persister cells to antibiotics while maintaining cells in an inactive state throughout a 24 h incubation period.

## Materials and methods

### Bacterial strains and growth conditions

The clinical isolate *S. aureus* SAU060112 (DSM 110939) and the type strain *E. coli* K12 (DSM 498) were used in this study. The strains were cultivated on tryptic soy agar, inoculated from single colonies into tryptic soy broth (TSB) in Erlenmeyer flasks and grown overnight at 37°C and at 180 rpm. The overnight cultures were diluted 1:1000 in TSB and incubated overnight again at 37°C and 180 rpm. This two-step inoculation was performed before each experiment. A modified version of M9 minimal salts (mM9) was used under starvation conditions to provide bacteria with salts, vitamins and metals for survival in the absence of a carbon source^28^. mM9 contains 1 × M9 salts (Sigma Aldrich), 2 mM MgSO_4_ (Sigma Aldrich), 0.1 mM CaCl_2_ (Sigma Aldrich), 1 mM Thiamine-HCl (Sigma Aldrich), 0.05 mM Nicotinamide (Sigma Aldrich), trace metals^5^.

### Minimum Inhibitory Concentration determination

The minimum inhibitory concentration (MIC) of ciprofloxacin and mitomycin C was determined using broth dilution in accordance with EUCAST regulations. Briefly, an overnight culture of *S. aureus* SAU060112 was added to 2-fold serially diluted antibiotics in TSB in 96-well plates to obtain a starting concentration of 5 × 10^5^ CFU/mL. 96-well plates were incubated at 50 rpm, 37°C overnight. Afterwards, MIC was determined as the minimum concentration where growth was inhibited by visually inspecting wells.

### Time-kill curves of exponential and stationary cultures in TSB or mM9 treated with ciprofloxacin or mitomycin C

For the exponential phase cultures, an overnight culture of *S. aureus* or *E. coli* was diluted 1:100 in TSB and incubated at 180 rpm, 37°C until reaching a turbidity 0.1-0.5. For the stationary phase cultures, the overnight culture was diluted from a turbidity of ∼ 10 to 1 in mM9 or TSB. The exponential or stationary cultures were then washed by centrifugation at 13150 × *g* for 10 min and the pellet was resuspended in the same volume fresh TSB or mM9. The washed culture was then diluted 10-fold into 1 mL TSB or mM9 with 10 ×, 50 × and 100 × MIC ciprofloxacin (Sigma Aldrich) or 10 × MIC mitomycin C (Sigma Aldrich), and incubated at 37°C, 50 rpm.

The CFU concentration at T_0_ was determined from the culture prior to mixing with antibiotics, and subsequently at set time-points by removing 100 μL from the samples, washing the cells in mM9 by adding 900 μL mM9 to the tube, centrifuging at 13150 × *g* for 10 min, and resuspending the pellet in 100 μL mM9. CFU was then determined by 10-fold dilution series and spotting out 10 μL (*S. aureus*) or 100 μL (*E. coli*) from each dilution step onto TSB agar.

### Rifampicin pre-treatment to induce persistence

Overnight cultures of *S. aureus* or *E. coli* was diluted to a turbidity of 1 in mM9 or TSB, washed by centrifugation as described above, resuspended in mM9 or TSB containing 100 μg/mL rifampicin and incubated for 30 min at 37°C, 180 rpm. Subsequently, the rifampicin-treated samples were washed by centrifugation and transferred to TSB or mM9 with ciprofloxacin as described above.

### Compound screening

Approximately 250 fragment compounds were purchased from Key Organics (Camelford, Cornwall) fragment library and screened resulting in a cluster of four fragments (compounds 241, 242, 243 and 244) having anti-persister activity and the same 3,5-disubstituted phenol motif. Compounds that incorporated this or a similar motif were purchased from Enamine Ltd (Kyiv, Ukraine). Compounds 258, 260, 267, 280, 292, 296 and 322 correspond to Enamine compound numbers Z2583036198, Z2668835953, Z2177044396, Z2683009586, Z1669437412, Z2683009578, Z4562301868 respectively. Over 400 compounds have been tested demonstrating the high throughput nature of this assay. Compounds were stored at room temperature until they were dissolved in DMSO at a concentration of 200 mM, at which point the compounds were stored at - 20°CAn overnight culture of *S. aureus* SAU060112 was diluted to a turbidity of 1 in mM9, washed by centrifugation as described above, and resuspended in equal volume of mM9. Compound treatments were carried out in 96-well plates where the test compounds and bacteria were added to mM9 buffer to obtain working concentrations of 1 mM, 5 mM and 10 mM compound and 5 × 10^7^ CFU/mL. The 96-well plate was incubated at 37°C, 50 rpm for 24 h and 10 μL from each well was then spotted on TSB agar and incubated overnight at 37°C. CFU was not enumerated, but a rough assessment was made, based on the following: > 10^4^ CFU/mL resulted in confluent growth, 10^2^ -10^4^ CFU/mL resulted in distinct colonies, and <10^2^ CFU/mL resulted in no growth.

### Statistical analyses

Data was tested for normality using a Shapiro-Wilk test. Normally distributed data was tested using a t-test for single comparisons and a one-way ANOVA using mixed-effects analysis for time kill curves and ordinary ANOVA for bar graphs and death rates. Equal variability was not assumed and the Geisser-Greenhouse correction was used. A post-hoc uncorrected Fisher’s LSD was performed for one-way ANOVA tests. Non-normally distributed data was tested using a Mann-Whitney U-test and for matched comparisons a Wilcoxon matched-pairs signed rank test was run. For normally distributed data, the mean ± standard deviation is shown, whereas for non-normally distributed data the median ± [range] is shown. GraphPad Prism was used for all statistical analyses (v. 9.5.1 (733) for Windows, GraphPad Software, San Diego, California USA, www.graphpad.com).

## Results and discussion

In this study, we used ciprofloxacin, a fluoroquinolone antibiotic targeting DNA gyrase, as the standard antibiotic to benchmark the effect of novel compounds against persister cells. Mitomycin C was included as a positive control, due to its well documented antimicrobial effect on bacterial persister cells^29^. We first determined the MIC values for these antibiotics against *S. aureus* SAU060112 (Table 1).

**Table 1.** Minimum Inhibitory Concentrations of used antibiotics and bacterial strains.

| Organism | Ciprofloxacin ($\mu\text{g/mL}$) | Mitomycin C ($\mu\text{g/mL}$) |
| --- | --- | --- |
| *S. aureus* SAU060112 | 1 | 0.8 |
| *E. coli* K12 | 0.008 | NA |

We first sought to determine, whether the concentration of bacteria and antibiotics affected what we perceived as persister cells; i.e., the fraction of viable cells after antibiotic treatment. These parameters are important to standardize, as the antimicrobial efficacy of antibiotics can depend on cell concentration (i.e. the inoculum effect)^30,31^, and because the antibiotic concentration must exceed the level, where bacterial death can be induced by activation of prophages in the genome, rather than by the direct action of the antibiotic^32^. Prophages are common in the genomes of clinical isolates of *S. aureus* and are also present in SAU060112^33^.

A time-kill assay was performed in TSB using four different turbidities (0.001, 0.01, 0.1 and 1) of exponential phase *S. aureus* cultures (Fig. 1). We chose to quantify viable cells as CFU after incubation times of 3 h, 5 h and 24 h, as this is the antibiotic exposure time that most studies have used for quantification of persister cells^20,27,34^. At turbidities 0.001 and 0.01, the cell concentration was below the detection limit at 3 h and 5 h, respectively (Fig. 1A and 1B), and the fraction of surviving cells could therefore not be determined. At turbidities 0.1 and 1, we measured a 3-log reduction in response to ciprofloxacin treatment. Therefore, we decided to use the turbidity of 0.1 (10^7^ CFU/mL) as a starting concentration for the remainder of the study.

**Figure 1.**
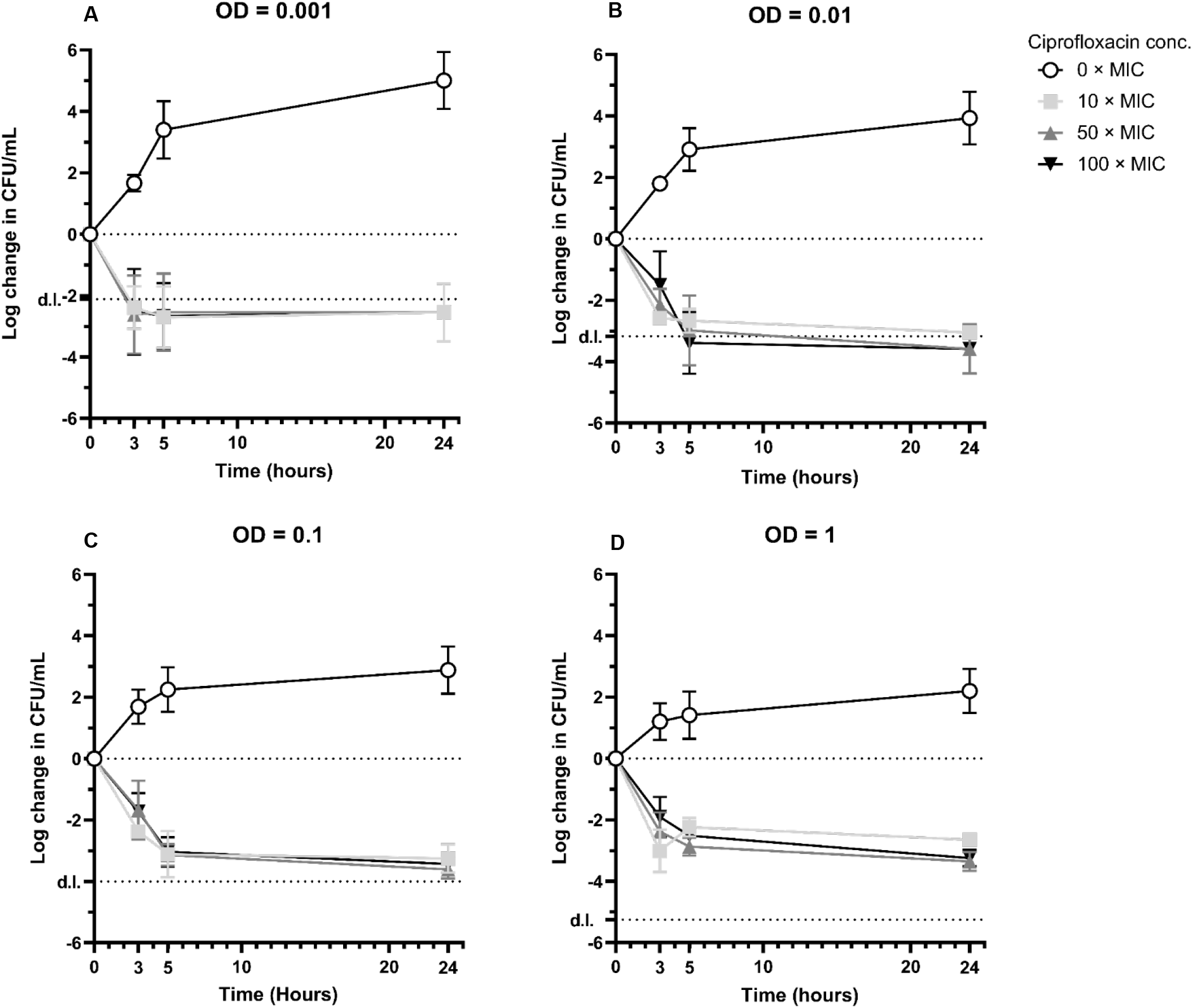
The effect ciprofloxacin concentration and bacterial concentration on time-kill curves and fraction of persister cells. An exponential phase cultures of *S. aureus* was washed and resuspended in TSB at the following optical densities of 1 (**A**), 0.1 (**B**), 0.01 (**C**) and 0.001 (**D**) in TSB before being treated with different concentrations of ciprofloxacin at time = 0 h. At 0 h, 3 h, 5 h, and 24 h viable cells were quantified as CFU. n ≥ 3, d.l.: detection limit.

### Rifampicin pre-treatment induced persister phenotype in *S. aureus* for at least 5 h

The fraction of persister cells is highly dependent on their metabolic state, and we therefore compared the fraction of persister cells in exponential- and stationary-phase cultures, which were exposed to 10, 50 or 100 × MIC of ciprofloxacin for 3, 5 or 24 h. Exponential phase cultures displayed the characteristic biphasic time-kill curve (Fig. 2A), and plateaued after 5 h treatment with no further reduction in CFU (one-way ANOVA, p > 0.05). Only 0.048 % ± 0.033 of the cells remained viable after 24 h treatment, which is similar to the fraction of persister cells reported by others^20^. This fraction of persister cells is too low to study a further decline in viable cells during subsequent screening for novel antibiotics with activity against this population. We therefore proceeded to quantify antibiotic tolerance in stationary phase cultures, which have been reported to contain a higher fraction of persister cells due to activation of various stress responses that are associated with persistence^35–37^.

**Figure 2.**
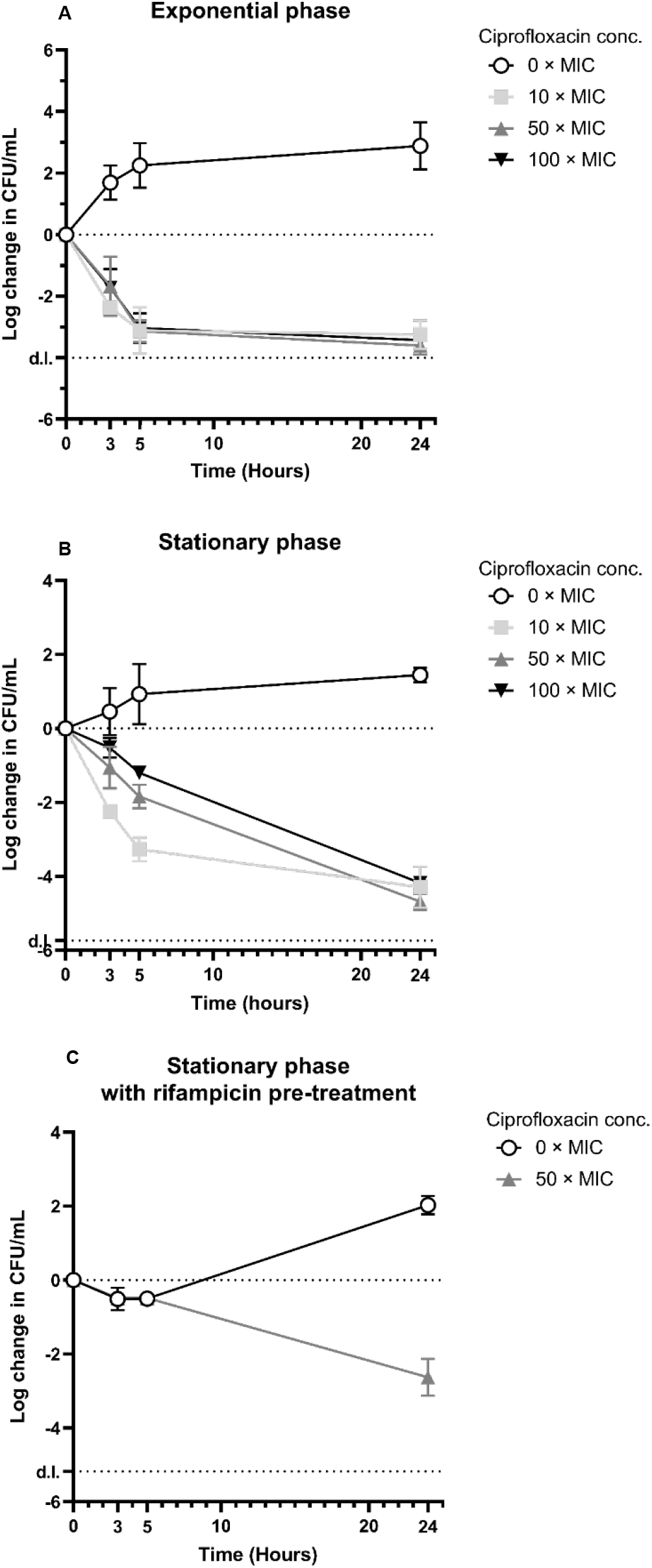
Time-kill curves of *S. aureus* in TSB. An exponential phase culture **(A)** or a stationary phase culture **(B)** was transferred to fresh TSB with or without ciprofloxacin at T = 0 h (1 × MIC = 1 μg/mL). **(C)** A stationary phase culture of S. aureus was transferred to fresh TSB with 100 μg/mL rifampicin for 30 min and subsequently washed and transferred to fresh TSB with or without ciprofloxacin at T = 0 h. Detection limit: d.l., n ≥ 3

The death rate was much slower for stationary phase compared to exponential phase cultures at the two highest ciprofloxacin concentrations (At 50 × MIC: -0.181 ± 0.018 vs -0.615 ± 0.061 log(CFU)·h^-1^, one-way ANOVA p < 0.0001; at 100 × MIC: -0.171 ± 0.012 vs -0.567 ± 0.102 log(CFU)· h^-1^, one-way ANOVA, p < 0.0001). However, at 10 × MIC the initial death rate was similar in stationary- and exponential phase cultures (−0.684 ± 0.069 vs -0.695 ± 0.138 log(CFU)·h^-1^, one-way ANOVA, p = 0.8426) and this sample was the only stationary phase culture that displayed a biphasic time-kill curve (Fig 2B). The fast initial death rate at the lowest ciprofloxacin concentration could indicate activation of prophages in the genome^22,38^. Earlier studies have shown that low antibiotic concentrations lead to prophage activation and thereby a higher killing rate compared to higher antibiotic concentrations. It is therefore important to use antibiotic concentrations that are high enough to avoid this effect.

The median fraction and range of stationary phase cells surviving 24-hour treatment at all ciprofloxacin concentrations was 0.0037% ± [0.0011,0.0153], which, unexpectedly, was lower than for exponential-phase cells (Mann-Whitney U-test, p < 0.0001). What would have appeared as increased tolerance in stationary phase cultures sampled after 3 or 5 h antibiotic exposure was thus reversed after 24 h exposure. It is important to note that the death rate of cells transferred from stationary phase cultures to TSB with antibiotics reflects both the antimicrobial action of the drug on non-growing stationary phase cells, but also the antimicrobial action on cells that are resume growth and loose the tolerant phenotype as nutrients become available. The death rate observed for stationary phase cultures may thus simply reflect the rate at which bacteria resume activity after transfer to TSB, rather than the rate at which ciprofloxacin kills non-growing cells. The rate with which bacteria resume activity after transfer from nutrient-poor to nutrient-rich conditions depends on the mechanisms the cells use to resume metabolic activity and protein synthesis. For example, Gram negative *E. coli* must reactivate dimerized ribosomes before protein synthesis can resume^34^. Cells in stationary phase quickly resume growth after transfer to fresh media, and inducing a non-growing state through other means than lack of nutrients could perhaps delay or slow down the rate of reactivation after transfer to TSB. In *S. aureus*, dimerized ribosomes do not disassemble easily upon transfer to fresh media and dimerization occurs in all growth phases, in contrast to *E. coli*^39^. Although the role of dimerized ribosomes in *S. aureus* has not been fully elucidated, there are thus contrasts in the proces between species that may affect how persister cells form.

Others have induced the persister phenotype in *E. coli* by inactivating the cells with antibiotics, using e.g. a short 30 min exposure to rifampicin, which stops transcription. This led to a ∼1,000 to ∼10,000-fold increase in tolerance to 5 μg/mL ciprofloxacin with 59% of the population displaying the persister phenotype after 3 h of ciprofloxacin treatment^20^. We therefore proceeded to determine if the same could be achieved for *S. aureus*. Rifampicin pre-treatment induced a non-dividing state, which lasted at least 5 h after transferring the culture to TSB (Fig. 2C). There was an initial loss (∼66%) of cells as a consequence of rifampicin exposure, but all the surviving cells displayed the persister phenotype and tolerated the subsequent ciprofloxacin treatment for at least 5 h. Hence a short rifampicin treatment protected the bacteria against ciprofloxacin, which corroborates similar results obtained with *E. coli*^20,24^. However, at incubation times > 5 h, the cells resumed activity, resulting in bacterial growth in the untreated controls, and bacterial killing in samples with ciprofloxacin. After 24 h incubation, 0.38 % ± 0.31 of cells had survived exposure to 50 × MIC ciprofloxacin. In conclusion, rifampicin pre-treatment can be used to generate a high concentration of persister cells, but the induced persister state only lasts a few hours.

### Continued starvation keeps the full *S. aureus* population in a persister phenotype

Ideally, screening for novel antimicrobials against persister cells should use a culture with a high concentration of persister cells, such that a 1000-fold reduction in viable cells can be detected. Furthermore, the fraction of tolerant cells should remain constant during the incubation. To enable longer incubation times, we therefore investigated if bacteria could simply be kept in a starved state during antibiotic exposure. In order to keep cells starved, we transferred exponential phase or stationary phase cells into a modified minimal salt solution (mM9 buffer), which is a carbon-free minimal medium containing vitamins, trace metals, divalent metal ions and M9 salts.

The exponential phase culture transferred to mM9 buffer displayed an initial loss of CFU in all samples including the untreated control (Fig 3A). This is most likely due to the abrupt transfer from a fast-growing state in a nutrient-rich medium to a nutrient-free medium. The CFU concentration in the control then increased between the 3 h and 5 h sampling point, indicating that some cells had recovered from the transfer and could be detected as CFU. The death rate from 5-24 h was the same for all samples, including the untreated control (one-way ANOVA, p > 0.05). In summary, the number of surviving cells in the untreated sample was higher than the ciprofloxacin treated samples at the 5 h and 24 h timepoints, but since the death rate from 5-24 h incubation was identical in all samples, all cells in the population must be tolerant to ciprofloxacin from 5 h onwards. While only 0.11 % ± 0.10 of the population survived 24 h exposure to ciprofloxacin, this number corresponds to 30.2 % ± 11.9 of CFU in the untreated sample. We therefore conclude that this approach results in a population with a relatively high fraction of antibiotic-tolerant cells, but with a high “background” death rate caused by starvation.

**Figure 3.**
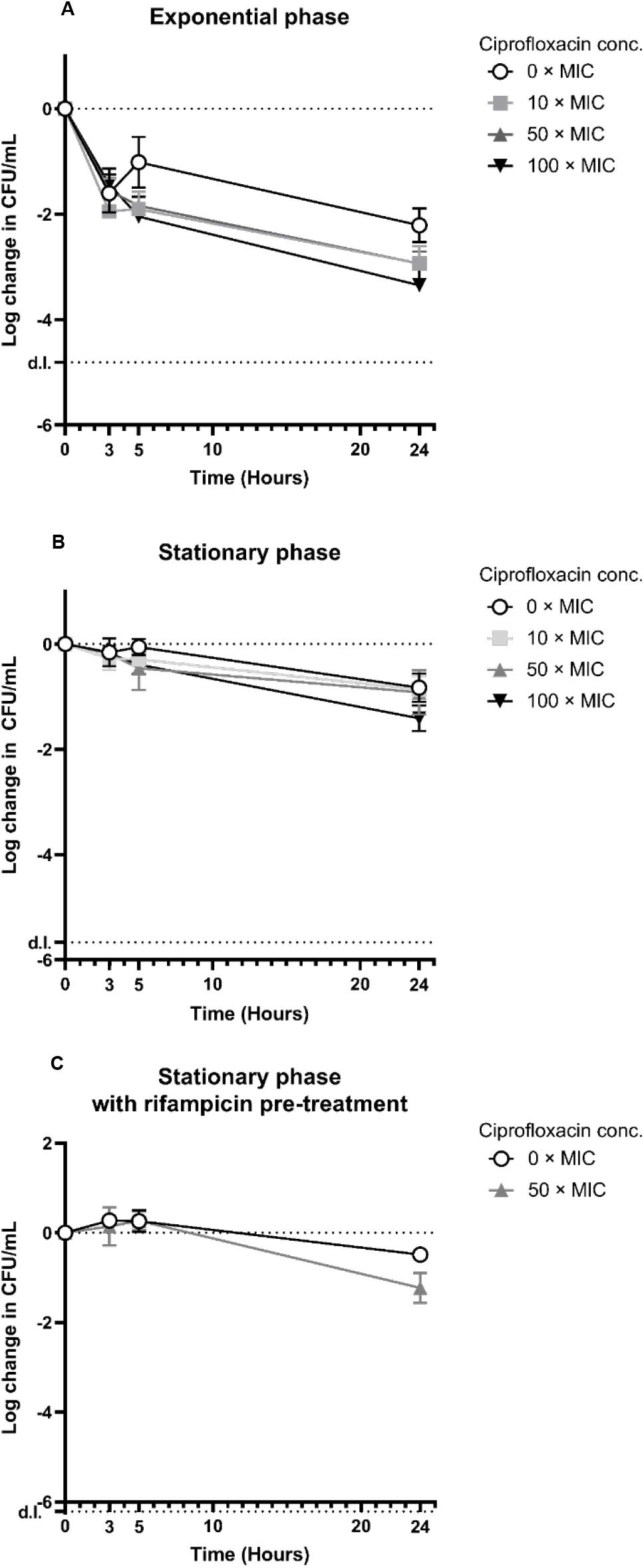
Time-kill curves of *S. aureus* in mM9 buffer. An exponential culture **(A)** or a stationary culture **(B)** grown in TSB was transferred to mM9 buffer with or without ciprofloxacin at 0 h (1XMIC = 1 μg/mL). **(C)** A stationary culture of *S. aureus* grown in TSB was transferred to mM9 buffer + 100 μg/mL rifampicin for 30 min and subsequently washed and transferred to mM9 buffer with or without ciprofloxacin at T = 0 h. Detection limit: d.l., n ≥ 3

We hypothesized that stationary phase cells would be better equipped to handle the transition from TSB to mM9, as metabolic processes have slowed down and mechanisms to cope with starvation have been activated. We therefore repeated the experiment with stationary phase cultures (Fig. 3B). In this case, the death rate was lower than for exponential phase cultures and 17.0 ± 10.2 % of the untreated control population remained viable after 24 h. Ciprofloxacin had no effect on viability at 3, 5 and 24 h when treating at 10 × or 50 × MIC (Mann-Whitney *U*-test, p > 0.05) and only a small effect was seen after 24 h at 100 × MIC (p = 0.0286) (Fig. 3B), indicating that all cells in the population were tolerant to antibiotics at the onset of the incubation. The time-kill curve did not display the biphasic shape, which was also not expected, given that the culture did not consist of two sub-populations with high and low antibiotic tolerance, and that there was no opportunity for cells to resume growth after transfer to the mM9 buffer. These results therefore demonstrate that transferring a stationary phase culture to a carbon-free minimal medium leads to a high survival rate, and 100% of the cells display the persister phenotype when challenged with fluoroquinolone antibiotics.

Although antibiotic exposure under starvation provided the antibiotic tolerance we had aimed for, we investigated if the persister-inducing treatment with rifampicin prior to incubation with ciprofloxacin affected the overall survival and tolerance to ciprofloxacin under these conditions. Again, we observed no significant difference in CFU between untreated and ciprofloxacin-treated cells after 3 or 5 h incubation (3 h: t-test, p = 0.6242, 5 h: Mann-Whitney U-test, p > 0.9999) (Fig. 3C). However, ciprofloxacin did affect cell viability after 24 h at 50 × MIC, compared to the untreated control (t-test, p = 0.0086). We therefore chose to proceed with the assay using a stationary culture transferred to mM9 without pre-treating with rifampicin.

### Continued starvation only keeps a small fraction of *E. coli* in a persister phenotype

Due to species-to-species differences in growth and stress responses, we decided to investigate differences in tolerance between *S. aureus* and a ciprofloxacin-susceptible *E. coli* isolate. *E. coli* and *S. aureus* stationary phase cultures were divided into two groups. One group received 30 min rifampicin treatment to induce the persister phenotype, while the other group did not. All samples were then transferred to TSB or mM9 buffer containing 50 × MIC ciprofloxacin, and incubated for 24 h.

When stationary phase cultures held in starvation in mM9 buffer, 86.0 % ± 31.8 of the *S. aureus* population tolerated ciprofloxacin for 24 h (Fig. 3B and 4A), while only 0.29 % ± 0.27 of *E. coli* survived (relative to the untreated control) (Fig. 4B). Pre-treatment with rifampicin did not improve the antibiotic tolerance of *E. coli* in mM9 buffer (one-way ANOVA, p = 0.8816). Antibiotic treatment in TSB resulted very low survival rates of both *S. aureus* and *E. coli* (2.25×10^−3^ % ±1.02×10^−3^and 0.93×10^−3^ % ±0.26×10^−3^, respectively). Pre-treatment with rifampicin did increase the fraction of tolerant cells by several orders of magnitude, but not above the the level observed in mM9 buffer. This was expected, as the data in Fig. 3C showed that rifampicin’s effect on antibiotic tolerance did not last for the full 24 h incubation period.

**Figure 4.**
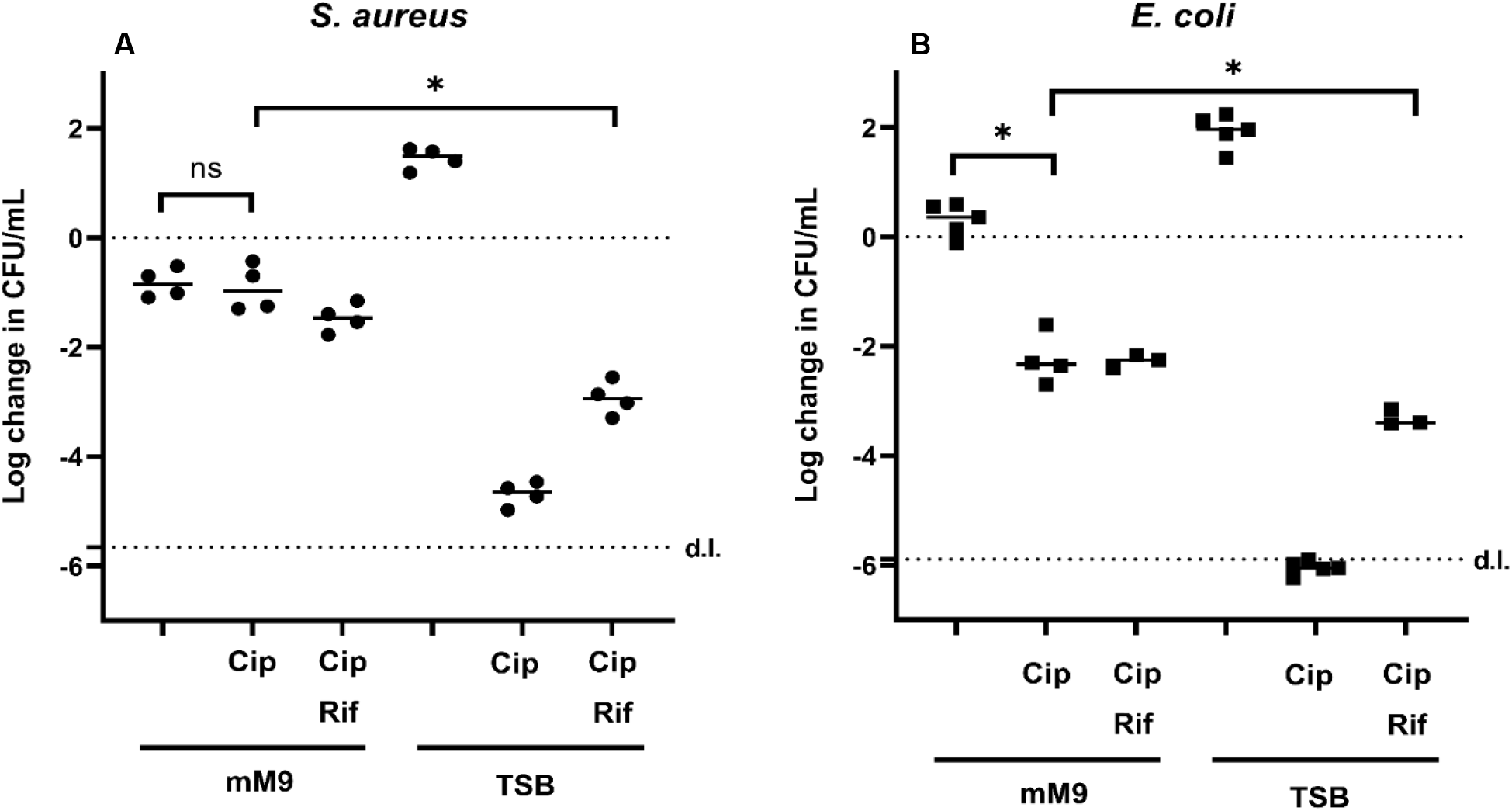
Change in log CFU/mL after 24 h treatment. *S. aureus* (A) or *E. coli* (B) was grown in TSB overnight and transferred to either TSB or mM9 buffer with or without 50 × MIC ciprofloxacin (Cip) (for *S. aureus* 1X MIC = 1 μg/mL, for *E. coli* 1X MIC = 8 ng/mL). Samples receiving 30 min 100 μg/mL rifampicin pre-treatment prior to ciprofloxacin treatment were included (Rif). *p < 0.0001, ns: p > 0.05, one-way ANOVA, n ≥ 3.

In summary, we conclude that controlled starvation obtained by transfer of stationary-phase cells to a carbon-free minimal medium can be used to generate a high concentration of bacteria, which displays the antibiotic tolerance of the persister phenotype over a 24 h incubation with antibiotics. This method is thus suitable for preparing bacteria for high-throughput screening to discover antimicrobials that target non-growing, antibiotic-tolerant bacteria. If the incubation period with antibiotics is shorter (< 5 h), a short pre-treatment with rifampicin prior to antibiotic exposure in growth media is also a powerful way of inducing temporary antibiotic tolerance, which was shown for *E. coli* previously^20^ and for *S. aureus* in this study (Fig. 3 and 4).

There is much discussion in the field about what characterizes persister cells, e.g. whether persister cells are truly dormant or retain some activity in order to avoid killing by antibiotics^3,21,40^. Here, we generate tolerant *S. aureus* that survive 24 h of 50 × MIC ciprofloxacin by transferring stationary-phase cells to a minimal medium where they cannot increase metabolic activity or resume growth. It was previously reported that to generate true *E. coli* persister cells using starvation, the cells must experience starvation for seven weeks^25,41^. One could thus argue, that starved *S. aureus* are not persister cells, but merely resting cells that are not dividing due to lack of nutrients, but otherwise maintain some level of activity that enables fast reactivation after transfer to a nutrient-rich medium. This is very likely the case. However, for their application in discovery of novel antibiotics to treat biofilm infections, the key questions are 1) whether these cells represent the antibiotic-tolerant phenotype of cells in biofilm infections, and 2) whether antimicrobials that display activity against these starved cells will also display the same activity against biofilm infections. The answers to both questions remain unknown. Nevertheless, we show that starved cells display the same tolerance to ciprofloxacin and susceptibility to Mitomycin C, as others have described for persister cells^29,42^. Importantly, the antibiotic tolerance was sustained during the 24 h incubation, and we could detect more than a 100,000-fold reduction in viable cells when treating with Mitomycin C, which makes the assay highly sensitive. In our subsequent quest to discover molecule structures with antimicrobial activity against *S. aureus* persister cells, we chose to incubate cells in starvation conditions to select compounds based on their direct antimicrobial activity on non-growing cells over 24 h.

### Screening tolerant *S. aureus* has the potential to identify novel anti-persister drugs

We started with a broad selection of compounds that were smaller than common antibiotics, but with structural similarities to previously published compounds with activity against persister cells^43^. Our strategy was to identify molecular structures in small molecules that have activity, such that these structures can be applied in design of more potent molecules with several active structures. This approach avoids screening compound libraries with large molecule structures that are unsuited for subsequent drug development.

We tested the antimicrobial activity of compounds at 1 mM, 5 mM and 10 mM, and subsequent rounds of compound design was then based on molecular structures that indicated activity. The antimicrobial efficacy was scored based on the absence of colonies (CFU/mL ≤10^2^), presence of discernible colonies (CFU/mL =10^2^-10^4^), or a confluent layer of bacteria (CFU/mL ≥10^4^) after spotting 10 μL of the bacterial suspension on agar. As the cell concentration in the untreated control was approximately 10^6^ CFU/mL, any effect detected by this assay would indicate at least a 2-log reduction in CFU. 50 × MIC mitomycin C and 50 × MIC ciprofloxacin were included as controls to verify the persister phenotype. Initial screening of approximately 250 compound fragments resulted in a cluster of four compounds having the same 3,5-disubstituted phenol motif (Fig. 5). Commercially available compounds having this or a similar structure to this active fragment were selected for testing. Of these tested compounds, compound 258 (Fig. 5) was identified as the most promising with antimicrobial activity resulting in >log 5 reduction in viable cells at 1 mM (Table 2). Further screening rounds of commercially available compounds found compounds with similar activities that had better physiochemical properties than compound 258, but none that were more active. In future compound design strategies, compounds will be based on 258 with focus on increasing the antimicrobial activity. For the compounds to be clinically relevant, activity should be in the micromolar range. None of the compounds so far have fulfilled this criterium, but the screening has yielded a potential for continuing to search for novel antibiotics.

**Table 2.** *S. aureus* persister cell reduction (Δlog) in response to compound treatment. *S. aureus* persister cells were treated with the numbered compounds of different concentrations. After 24 h treatment, each sample was spotted out on an agar plate and incubated overnight. The survivors were quantified from the agar plates in units of CFU/mL and normalised to the CFU/mL at the start of the incubation (10^7^). ≤ -5 denotes full inhibition/no growth since the detection limit of the assay is ≤10^2^ CFU/mL. ≥ -2 CFU/mL denotes no detectable inhibition/full growth. For any sample where there is inhibition, but no countable colonies, the log change of survivors is between -5 and -2 (denoted > -5, < -2). n = 3. NA: This compound concentration was not tested.

| Compound | 1 mM | 5 mM | 10 mM |
| --- | --- | --- | --- |
| 258 | $\leq -5$ | $\leq -5$ | $\leq -5$ |
| 260 | $\geq -2$ | $\leq -5$ | $\leq -5$ |
| 267 | $> -5, < -2$ | $> -5, < -2$ | $> -5, < -2$ |
| 280 | $> -5, < -2$ | $\leq -5$ | $\leq -5$ |
| 292 | $\geq -2$ | $> -5, < -2$ | $> -5, < -2$ |
| 296 | $> -5, < -2$ | $> -5, < -2$ | $> -5, < -2$ |
| 322 | $> -5, < -2$ | $\leq -5$ | $\leq -5$ |
| | 10 × MIC | 50 × MIC | 100 × MIC |
| Ciprofloxacin | $\geq -2$ | $\geq -2$ | $\geq -2$ |
| Mitomycin C | $\leq -5$ | $\leq -5$ | NA |

**Figure 5.**
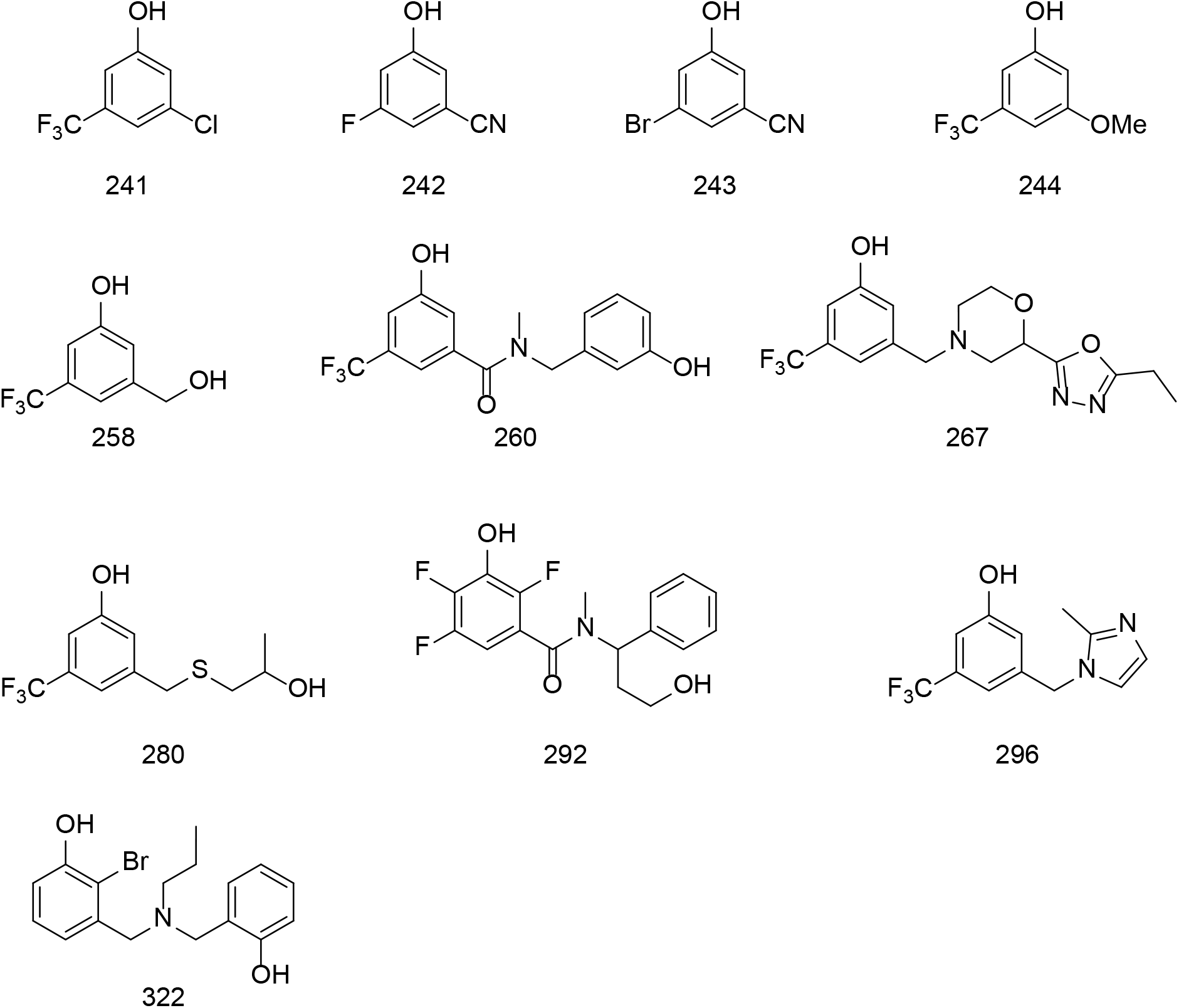
The structures of potential anti-persister compounds tested against starvation-induced tolerant cells. 250 compounds were pre-screened for anti-persister activity yielding a cluster of four fragments (compounds 241, 242, 243 and 244) with the same 3,5-disubstituted phenol motif. This motif was used to design potential anti-persister compounds that were further screened using the method demonstrated here.

### Strategies for high-throughput screening to identify anti-persister drugs

The key to high-throughput screening is scalability and automatization. In our assay, we assessed bacterial viability by scoring CFU on agar without prior dilution. An even simpler approach would be to transfer the 10 μL to TSB in a new microwell plate and score the outcome based on presence/absence of viable cells that would lead to growth measured as optical density during a subsequent incubation. This transfer and readout can be fully automated with common robotic systems but would also narrow the read-out to a yes/no answer to whether any viable cells remained after the incubation.

Other high-throughput assays have assessed viability based on loss of membrane permeability using membrane-impermeable fluorescent DNA-binding dyes. Using a fluorescent readout also enables automated analysis of the result in a microwell plate format, but the approach will be biased toward identifying membrane-active compounds, or compounds that lead to cell lysis through other means. A highly effective drug like Mitomycin C would be missed in this assay, as it kills cells by crosslinking DNA and does not immediately lead to cell lysis (data not shown). The approach was applied to a 85,000 compound library, leading to the identification of compound NH125, a bacterial histidine kinase inhibitor with anti-persister activity^44^. The same group used an MRSA killing screening method in the nematode *Caenorhabditis elegans* to screen 82,000 compounds and identified a new class of synthetic retinoids that are effective against MRSA persister cells^45^.

Some anti-persister compounds kill persister cells together with conventional antibiotics by “waking up” the persister cells, and thereby disarming defence mechanisms^46–48^. Screening for compounds with this effect must occur in a nutrient-rich medium and in combination with other antibiotics in order to identify the killing propensity of the combination. Otherwise, the compounds should be screened based on persister resuscitation and not persister killing, which likewise has been shown to be a successful approach^48^. Our assay is therefore not suited to identify this type of anti-persister drugs, but our results indicate that such drugs could be identified by inducing the persister phenotype through rifampicin exposure and subsequently testing the antimicrobial effect in TSB with ciprofloxacin during < 5h incubation.

One of the newest trends in compound screening is the use of machine learning with training and predictions to initiate a screening process. Stokes et al used deep learning of antibiotic discovery, where they used a training set of 10^4^ compounds to initiate a baseline for a successful compound^49^. Subsequently, they carried out a large-scale prediction of 10^8^ small molecules in a chemical landscape. When they had narrowed the compounds down to 10^5^-10^6^ compounds, they used a conventional molecule screening method for validating the hits by incubating *E. coli* in nutrient rich media containing the selected compounds to determine growth inhibition. This approach led to the discovery of halicin, a molecule that is structurally divergent from conventional antibiotics and displays bactericidal activity against active cells as well as inactive persister cells. The anti-persister activity of halicin was perhaps a stroke of luck since the discovery process was not targeted towards anti-persister compounds. However, the use of machine learning to identify bactericidal (rather than bacteriostatic) drugs increases the chance of finding antibiotics that kill persister cells. Machine learning can be very efficient when initialising a new screening of bactericidal drugs. However, it must always be followed with experimental evidence, and high-throughput assays that reliably detect anti-persister activity with high sensitivity are thus needed to aid the discovery of novel antibiotics to treat recalcitrant bacterial infections harbouring persister cells. With our method there is a potential to discover new anti-persister drugs by screening cells that are highly tolerant to conventional antibiotics.

## Acknowledgements

This work was supported by the Novo Nordisk Foundation (grant no. NNF19OC0058357).

